# RCytoGPS: An R Package for Reading and Visualizing Cytogenetics Data

**DOI:** 10.1101/2021.03.16.389791

**Authors:** Zachary B. Abrams, Dwayne G. Tally, Lynne V. Abruzzo, Kevin R. Coombes

## Abstract

**Summary:** Cytogenetics data, or karyotypes, are among the most common clinically used forms of genetic data. Karyotypes are stored as standardized text strings using the International System for Human Cytogenomic Nomenclature (ISCN). Historically, these data have not been used in large-scale computational analyses due to limitations in the ISCN text format and structure. Recently developed computational tools such as CytoGPS have enabled large-scale computational analyses of karyotypes. To further enable such analyses, we have now developed RCytoGPS, an R package that takes JSON files generated from CytoGPS.org and converts them into objects in R. This conversion facilitates the analysis and visualizations of karyotype data. In effect this tool streamlines the process of performing large-scale karyotype analyses, thus advancing the field of computational cytogenetic pathology.

**Availability and Implementation:** Freely available at https://CRAN.R-project.org/package=RCytoGPS

**Supplementary information:** Supplementary data are available at *Bioinformatics* online.

## 1 Introduction

Cytogenetic data are a commonly used form of genetic data, particularly in clinical pathology. Advancements in the treatment of many diseases have come from a better understanding of the underlying cytogenetic events related to that disease. One such example is the Philadelphia translocation and its relation to chronic myeloid leukemia (Johansson, et al., 2002). However large-scale processing of cytogenetic data has historically been difficult due to the format in which karyotype data are stored.

Karyotype data are stored in a text-based structure called the International System for Human Cytogenomic Nomenclature (ISCN) (McGowan-Jordan, et al., 2016). Although this standard is effective in recording cytogenetic abnormalities, it is not an inherently easy language for computers to read. Recently, new computational tools have been developed to parse ISCN karyotypes. One such tool is the CytoGPS online system, available at http://cytogps.org/ (Abrams, et al., 2020; Abrams, et al., 2019). CytoGPS works by breaking ISCN karyotypes into a three-part binary mapping structure termed the Loss-Gain-Fusion (LGF) model. CytoGPS enables users to input text-based ISCN karyotypes and receive a JSON file containing the parsed LGF model data.

However, the CytoGPS website only parses and maps cytogenetic data; it does not help users analyze, visualize, or interpret their data. This is a limitation of the current CytoGPS system since not all users will have the time, knowledge, and/or resources to develop tools that perform the downstream analyses required for their research. To solve this problem while improving the computational utility of CytoGPS, we developed the RCytoGPS R package.

## 2 Methods

RCytoGPS uses the output JSON file generated from CytoGPS.org as its input data. A user may input one or more JSON files at a time. For single files, the user provides the path to the directory and specifies the file name. For multiple files, the user simply puts all the JSON files they wish to analyze into a local directory, and then provides that path to RCytoGPS. Then RCytoGPS reads in each JSON file and constructs an R structure to organize the data.

Processing CytoGPS JSON files with the readLGF function in RCytoGPS results in a list containing five elements: source, raw, frequency, size, and CL. Source is a character vector that holds the names of the input JSON files. Raw is a list of lists, one per input file, containing the binary LGF matrix for all karyotypes and a status report flagging any karyotypes that failed processing because of syntax or other errors. Frequency is a data frame summarizing the frequency of losses, gains, or fusions at the level of cytogenetic bands. Size is the total number of processed clones, since each karyotype can contain multiple clones. CL is a description of the cytoband locations (chromosome name, start and end base pairs) in build GRCh38 of the human genome, along with standard names of chromosome arms and cytogenetic bands. This information is organized such that it is easily accessible to the user should they wish to perform any custom or novel analyses of their own design.

One of the key elements to RCytoGPS is its ability to generate different visualizations to help interpret results. These visualizations are based on the frequency of cytogenetic events in the LGF model. To illustrate, we have processed all 4,999 karyotypes from AML samples in the publicly available Mitelman Database of Chromosome Aberrations and Gene Fusions in Cancer (Mitelman, et al., 1991). Genome-wide barplots (**Figure 1A**) clearly identify the most frequent losses (orange) on chromosomes 5 and 7 and the most frequent gain (green) on chromosome 8. By zooming in to chromosome 17 (**Figure** 1B), we can see that roughly 10% of cases have a loss of the entire chromosome, The plot also suggests that another 5% may have a loss of 17p and a gain of 17q, probably from an isochromosome 17q. These visualizations can help researchers identify regions of interest that can later be explored more carefully using the binary LGF vectors. A more complete set of example visualizations is contained in the “gallery” vignette included in the R package.

**Figure 1:**
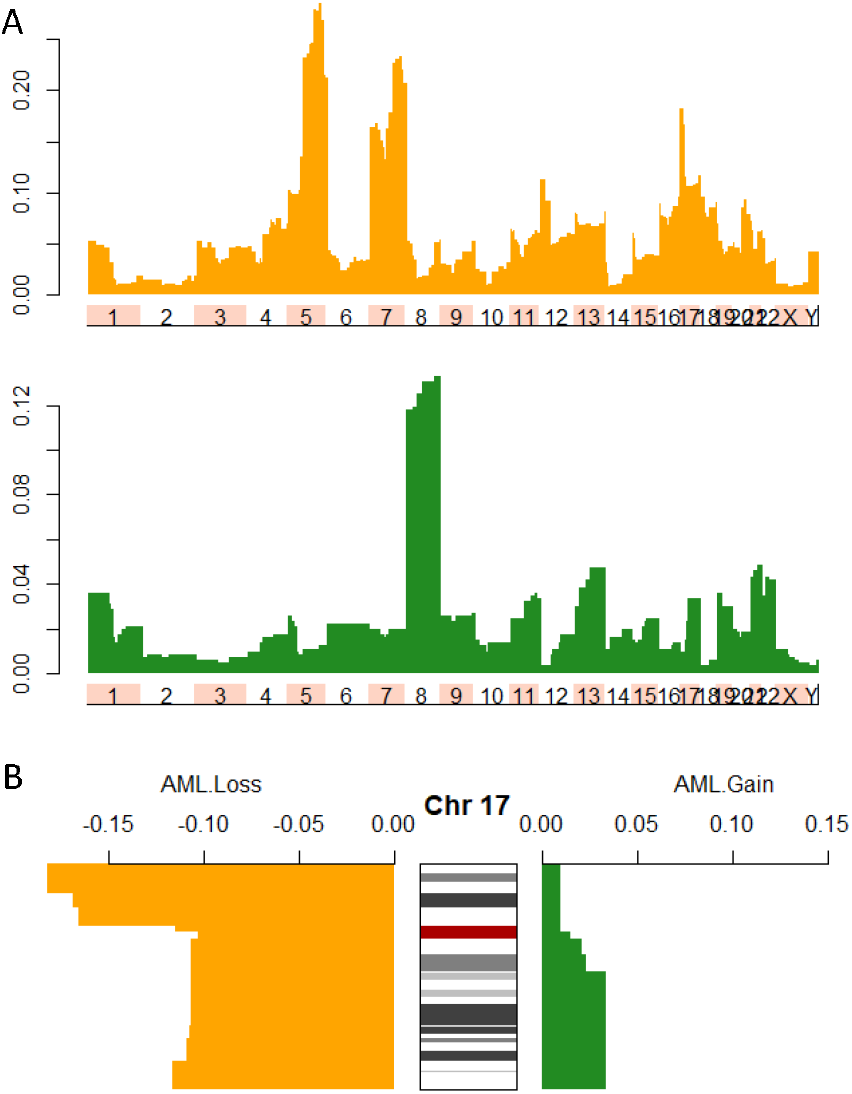
Visualizations of cytogenetic data of 4,999 AML karyotypes showing the frequency of losses (orange) and gains (green) in a dataset. (A) Barplots across the entire genome. (B) Barplots along chromosome 5, including visualization of cytobands.

Once the researcher has identified a potential chromosome of interest they can effectively zoom in on that region with a different set of visualizations. A single chromosome comparison is shown in Figure 1B. Here the loss and gain events are compared only along chromosome 5. The ideogram of the chromosome is shown in the middle along with the corresponding G banding pattern. The frequency of loss events is shown on the left side of the chromosome while gain events are shown on the right side. Users can choose whichever colors they wish to represent their data. This form of side-by-side comparison can also be used to isolate key differences between two cytogenetic data sets.

We note here that RCytoGPS does not provide tools for the further analysis nor visualization of the raw binary data matrices containing the LGF data. However, those matrices can serve as direct inputs to the Mercator R package, which supports clustering and visualization of binary data using a variety of distance metrics, clustering algorithms, and visualization tools (Abrams, et al., 2021)

## 3 Conclusion

RCytoGPS is an R package that helps researchers import and visualize cytogenetic frequency data after it has been processed by CytoGPS. By helping users easily transfer their cytogenetic data into R, storing the data in a user-friendly form, and providing an array of different visualization options, we hope that this tool can aid computational pathologists in their cytogenetic research.

